# Structure of population activity in primary motor cortex for single finger flexion and extension

**DOI:** 10.1101/2020.03.17.996124

**Authors:** Spencer A. Arbuckle, Jeff Weiler, Eric A. Kirk, Charles L. Rice, Marc Schieber, J. Andrew Pruszynski, Naveed Ejaz, Jörn Diedrichsen

## Abstract

How is the primary motor cortex (M1) organized to control fine finger movements? We investigated the population activity in M1 for single finger flexion and extension, using 7T functional magnetic resonance imaging (fMRI) in female and male human participants, and compared these results to the neural spiking patterns recorded in two male monkeys performing the identical task. fMRI activity patterns were distinct for movements of different fingers, but quite similar for flexion and extension of the same finger. In contrast, spiking patterns in monkeys were quite distinct for both fingers and directions, similar to what was found for muscular activity patterns. The discrepancy between fMRI and electrophysiological measurements can be explained by two (non-mutually exclusive) characteristics of the organization of finger flexion and extension movements. Given that fMRI reflects predominantly input and recurrent activity, the results can be explained by an architecture in which neural populations that control flexion or extension of the same finger produce distinct outputs, but interact tightly with each other and receive similar inputs. Additionally, neurons tuned to different movement directions for the same finger (or combination of fingers) may cluster closely together, while neurons that control different finger combinations may be more spatially separated. When measuring this organization with fMRI at a coarse spatial scale, the activity patterns for flexion and extension of the same finger would appear very similar. Overall, we suggest that the discrepancy between fMRI and electrophysiological measurements provides new insights into the general organization of fine finger movements in M1.

**Significance statement:** The primary motor cortex (M1) is important for producing individuated finger movements. Recent evidence shows that movements that commonly co-occur are associated with more similar activity patterns in M1. Flexion and extension of the same finger, which never co-occur, should therefore be associated with distinct representations. However, using carefully controlled experiments and multivariate analyses, we demonstrate that human fMRI activity patterns for flexion or extension of the same finger are highly similar. In contrast, spiking patterns measured in monkey M1 are clearly distinct. This suggests that populations controlling opposite movements of the same finger, while producing distinct outputs, may cluster together and share inputs and local processing. These results provide testable hypotheses about the organization of hand control in M1.

## Introduction

Dexterous movements of fingers require accurate coordination of different hand muscles. Hand muscles are innervated by motorneurons in the ventral horn of the spinal cord, which receive direct and indirect projections from the hand region of the contralateral primary motor cortex (M1) (Lemon, 2008). In monkey species capable of better finger individuation, direct (monosynaptic) projections from M1 to ventral horn motor neurons are more pronounced (Heffner & Masterton, 1983; Bortoff & Strick, 1993). Lesions to the corticospinal tract (Tower, 1940; Lawrence & Kuypers, 1968; Lawrence & Hopkins, 1976; Sasaki et al., 2004) or to M1 (permanent: Liu & Rouiller, 1999; Darling et al., 2009; reversible: Schieber & Poliakov, 1998) result in a significant loss of finger individuation. Such symptoms are also reported in human stroke patients who have damage to the hand area of M1 or the descending corticospinal pathway (Lang & Schieber, 2003; Xu et al., 2017). These results indicate that M1 is important for the fine control of individuated finger movements.

What is less well understood is how this cortical control module for finger movements is organized. Here, we studied this question by investigating cortical activation patterns evoked during flexion and extension of individual fingers. Previous electrophysiological work in macaque monkeys (Schieber & Hibbard, 1993; Schieber & Poliakov, 1998) have indicated that motor cortical neurons have complex tuning functions, often responding to movements of multiple fingers and to both flexion and extension movements. Therefore, there exists no clearly organized “map”, with separate regions dedicated to the control of a single finger. Instead, the population of M1 neurons involved in hand control must be organized by some other principle.

One plausible principle is that the statistics of natural hand use shapes the organization of neuronal populations in the hand region of M1. This idea predicts that movements that commonly co-occur in every-day life are represented in overlapping substrates in M1 (Graziano & Aflalo, 2007). In humans, fingers with high correlations between their joint-angle velocities during every-day hand movements (Ingram, et al., 2008) have been shown to have more similar M1 activity patterns, as measured with fMRI (Ejaz et al., 2015). The correlation structure of every-day finger movements nearly fully explained the relative similarities of M1 finger activity patterns, and fit the data better than a model that used the similarity of the required muscle activity patterns (i.e. predicting that movements that use similar muscles also have similar activity patterns) or a somatotopic model (i.e. predicting that fingers are represented in an orderly finger map).

In this paper, we asked to what degree this kinematic hypothesis could generalize to movements of the same finger in different directions. We measured the activity evoked in the hand area of M1 using high-field fMRI while human participants performed near-isometric single finger flexion and extension presses with their right hand. By extrapolating the model used in Ejaz et al. (2015) to this situation, we predicted that each movement should have its own, clearly separated representation in M1, as flexion and extension movements of the same finger can never co-occur. Indeed, it has been recently suggested that human motor cortex has multiple representations of each finger, one dedicated to flexion and one to extension (Huber et al., 2020).

We found, however, that the measured M1 fMRI patterns for flexion and extension of the same finger were strikingly similar, much more similar than would be expected for two movements that cannot co-occur. This similarity was not the result of co-contraction during the task. To better understand these results, we investigated the representational structure of single-neuron activity in M1 of two macaque monkeys trained on the same flexion-extension task (data from Schieber & Rivlis, 2005; Schieber & Rivlis, 2007). The spiking patterns in monkeys were quite distinct for fingers and directions. From these results, we propose two, non-mutually exclusive hypotheses about the organization of finger movement representations in the primary motor cortex.

## Materials and Methods

### Human participants

Nine healthy, participants were recruited for the study (5 males and 4 females, mean age=24.78, SD=4.68; mean Edinburgh handedness score=90.11, SD=11.34). Participants completed 3 experimental sessions. During the first training session, participants learned to perform the finger individuation task. In the scanner session, participants performed the finger individuation task while undergoing fMRI. In the EMG session, participants performed the finger individuation task while muscle activities were recorded. All participants provided informed consent before the beginning of the study, and all procedures were approved by the Office for Research and Ethics at the University of Western Ontario.

### Experimental design of human finger individuation task

In all three (training, scanning, and EMG) sessions, the five fingers of the right hand were individually clamped between two keys (Fig. 1A). Foam padding on each key ensured each finger was comfortably restrained. Force transducers (Honeywell-FS series, dynamic range=0-16N, resolution<0.02N, sampling rate=200Hz) above and below each key monitored the forces applied by each finger in extension and flexion directions.

**Figure 1.**
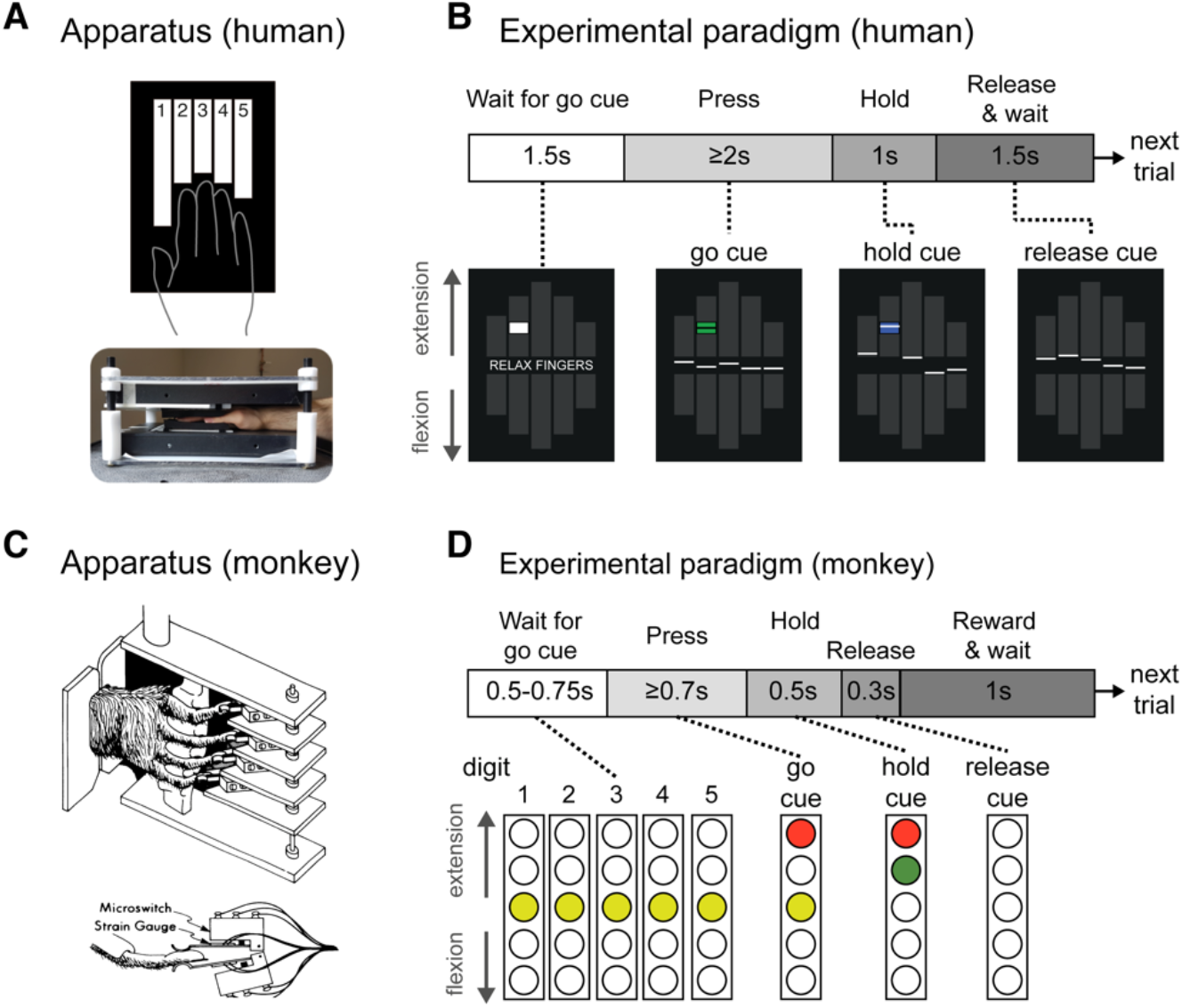
Experiment paradigms. (**A**) Human participants made isometric single finger presses in the flexion and extension directions on a custom-built keyboard. Each finger of the right hand was clamped between two keys, and each key was associated with a force transducer either above (keyboard on top of hand) or below (keyboard under the hand) the key to monitor forces applied in the flexion and extension directions, respectively. (**B**) Schematic illustration fo a single trial in the fMRI and EMG sessions, with associated visual feedback shown below. The white lines represent the produced force for each finger. Applying flexion to a finger key moved the associated line down (vice-versa for extension). The cue box (centred at target force) was initially presented as white at the trial start, and turned green to cue the participant to make the finger press (here, index finger extension). The box turned blue to instruct participants to maintain the current force. At the end of the press hold, the cue box disappeared and participants relaxed their hand. (**C**) The monkey hand configuration and device (illustration from Schieber, 1991). (**D**) Trial schematic for the monkey task. The columns represent 5 LED cues (one per finger) which instructed the monkey both what finger and what direction to press. The monkeys had up to 700ms from the onset of the go cue to press the cued finger in the cued direction. They were trained to hold the press for 500ms before relaxing the finger.

During the task, participants viewed a screen that presented two rows of five bars (Fig. 1B). These bars corresponded to flexion or extension direction for each of the five fingers of the right hand. The forces applied by each finger were indicated on the visual display as five solid white lines (one per finger). On each trial, participants were cued to make an isometric, single-finger flexion or extension press at one of three forces levels (1, 1.5, or 2N for extension; 1.5, 2, or 2.5N for flexion) through the display of a white target box (Fig. 1B). Extension forces were chosen to be lower than flexion forces, as extension finger presses are more difficult (Valero-Cuevas, Zajac, & Burgar, 1998; Li, et al., 2003) and can lead to more enslaving (i.e. co-articulation) of non-instructed fingers (Yu, Duinen, & Gandevia, 2010). This design yielded two levels of matched target forces for flexion and extension presses (1.5 and 2N). The forces were similar to the low forces required in the monkey task design. The finger displacement required to achieve these force thresholds was minimal, such that the finger presses were close to isometric.

Each trial lasted 6000ms and consisted of four phases (Fig. 1B): a cue phase (1500ms), a press phase (2000ms), a hold phase (1000ms), and a 1500ms inter-trial interval. This trial structure was designed to mirror the NHP task (see NHP methods and also Schieber, 1991). During the cue phase, a white box appeared in one of the ten finger bars presented on screen, indicating the desired finger and direction. The desired pressing force was reflected by the relative location of the cue within the finger bar. After 1500ms, the cue turned green. This instructed the participant to initiate the finger press. Participants had up to 2000ms after the cue turned green to reach the specified force. Once the pressing force was within the target box (target force ±12.5%) the cue turned blue. Participants were trained to hold the force constant within this interval for 1000ms. When this time had elapsed, the cue disappeared and the participants were instructed to release the press by relaxing their hand. Importantly, participants were instructed not to actively move the finger in the opposite direction. A new trial started every 6s. For the scanning session, periods of rest were randomly intermixed between trials (see below). The muscle recording sessions lacked these rest periods, but otherwise had the same trial structure.

Trials of the 30 conditions (5 fingers × 2 directions × 3 forces) were presented in a pseudo-random order. Trials were marked as errors if the participant was too slow (i.e. did not initiate movement within 2000ms of the go-cue), pressed the wrong finger or in the wrong direction, or if the participant did not reach at least 0.5N force with the cued finger in the cued direction. Due to the pre-training, the participants had low error rates in both the fMRI (mean error rate across conditions=1.48% ±1.05% sem) and EMG (mean error rate across conditions=1.30% ±0.97%) sessions, and accurately produced the required target forces (fMRI: mean peak force accuracy=108.93% ±2.56% of the target forces; EMG: mean accuracy=107.80% ±2.19%). Therefore, we included all trials in subsequent analyses.

We also did not exclude any trials based on finger co-activation. Overall, participants were able to individuate their fingers relatively well. During fMRI extension trials, the forces applied through the non-instructed fingers were, on average, 14.01% (±1.41%) of the forces applied by the instructed finger. During fMRI flexion, forces produced by non-instructed fingers was 20.51% (±1.49%) of the force produce by the instructed finger. Most enslaving occurred during presses of the middle, fourth, and little fingers, all of which are difficult to individuate (Schieber, 1991). Note, however, that the presence of enslaving does not compromise the main finding of our paper. To some degree, neural activity patterns related to flexion and extension of single fingers will always depend on the biomechanical coupling between fingers, either because the cortical activation patterns need to overcome that coupling, or because coupling does occur, which then influences the recurrent sensory input. Our main conclusions are based on comparisons between flexion and extension presses, and remain valid whether we study the actions of isolated fingers, or groups of fingers (see discussion).

### fMRI acquisition and analysis

#### Image acquisition

We used high-field functional magnetic resonance imaging (fMRI, Siemens 7T Magnetom with a 32 channel head coil at Western University, London, Ontario, Canada) to measure the blood-oxygen-level dependent (BOLD) responses in human participants. For each participant, evoked-BOLD responses were measured for isometric, single-finger presses in the flexion and extension directions.

There were 2 repeats of each condition during each imaging run (5 fingers × 2 directions × 3 force levels × 2 repeats = 60 trials). Trial order in each run was randomized. In addition, 5 rest conditions of 6000ms were randomly interspersed between trials within each run. Each run lasted approximately 390 seconds. Participants performed 8 such runs during the scanning session.

During each run, 270 functional images were obtained using a multiband 2D-echoplanar imaging sequence (GRAPPA, in-plane acceleration factor=2, multi-band factor=2, repetition time [TR]=1500ms, echo time [TE]=20ms, flip angle [FA]=45 deg). Per image, we acquired 32 interleaved slices (without gap) with isotropic voxel size of 1.5mm. The first 2 images in the sequence were discarded to allow magnetization to reach equilibrium. To estimate magnetic field inhomogeneities, we acquired a gradient echo field map at the end of the scanning session. Finally, a T1-weighted anatomical scan was obtained using a magnetization-prepared rapid gradient echo sequence (MPRAGE) with a voxel size of 0.75mm isotropic (3D gradient echo sequence, TR=6000ms, 208 volumes).

#### Image preprocessing and first-level analysis

Functional images were first realigned to correct for head motion during the scanning session (3 translations: x,y,z; 3 rotations: pitch, roll, yaw), and co-registered to each participant’s anatomical T1-image. Within this process, we used a B0 fieldmap to correct for image distortions arising from magnetic field inhomogeneities (Hutton et al., 2002). Due to the relatively short TR (1.5s), no slice-timing correction was applied. Nor was the data spatially smoothed or normalized to a standard template.

The minimally preprocessed data were then analyzed using a general linear model (GLM; Friston et al., 1994) using SPM12 (fil.ion.ucl.ac.uk/spm/). Each of the finger-direction-force conditions were modeled with separate regressors per run, resulting in 30 regressors per run (30*8 runs = 320 task regressors), along with an intercept for each run. The regressor was a boxcar function that started at the presentation of the go-cue and lasted for the trial duration, spanning the press, hold, and release periods of each trial. The boxcar functions were convolved with a hemodynamic response function with a delayed onset of 1000ms and a post-stimulus undershoot at 7500ms. Given the low error rate, we did not exclude any trials from this analysis. To model the long-range temporal autocorrelations in the functional timeseries, we used the SPM FAST autocorrelation model with restricted-maximum likelihood estimation (see Arbuckle et al., 2019 for details). High-pass filtering was then achieved by temporally pre-whitening the functional data with this temporal autocorrelation estimate. This analysis resulted in one activation estimate (“beta-weights”) for each of the 30 conditions per run for each participant. For visual display (as in Figure 2) and further analysis, the beta values were divided by the root-mean-square error from the first-level GLM to yield a *t*-value per voxel for each condition in each run.

**Figure 2.**
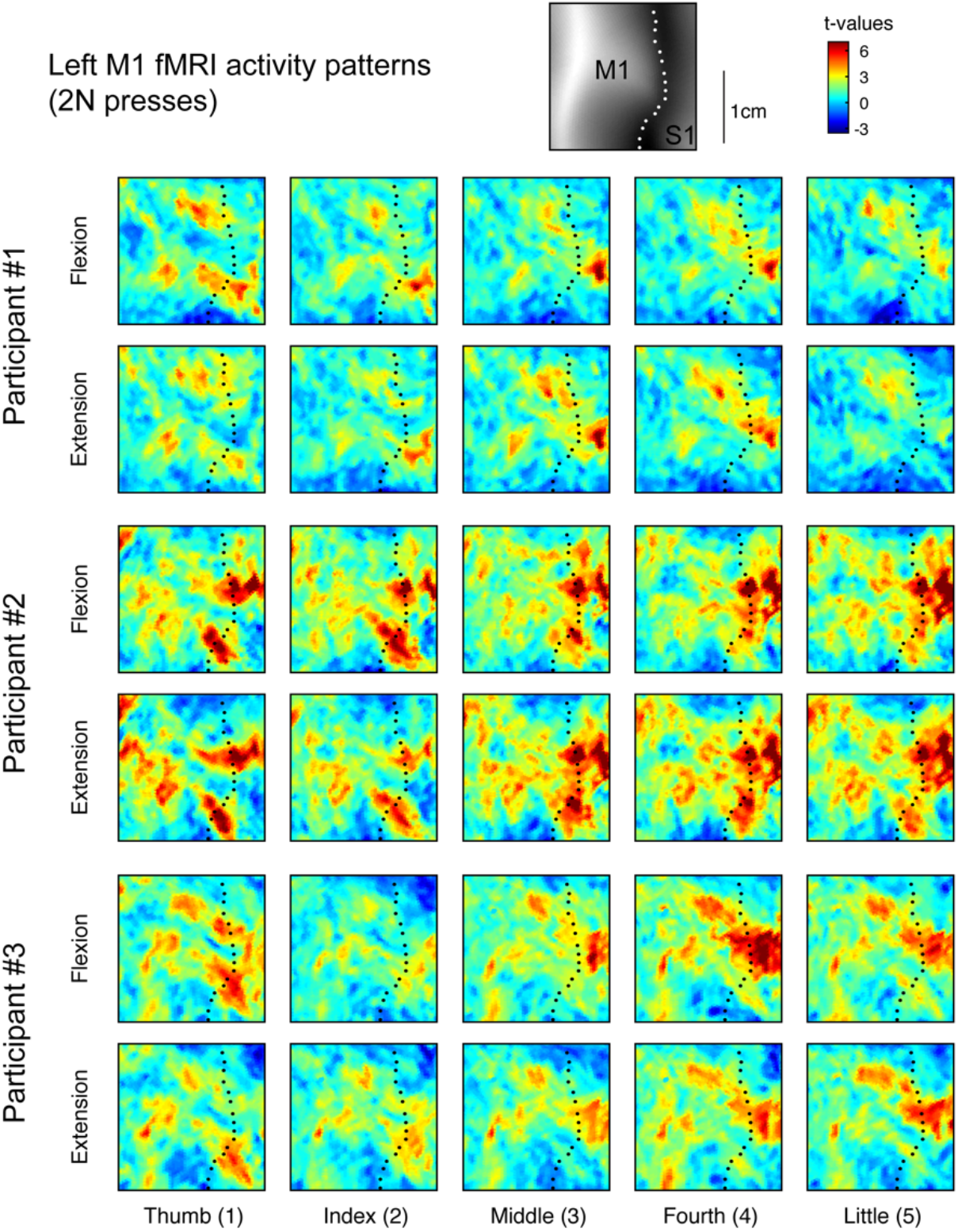
fMRI activity patterns for finger flexion and extension in human M1. Evoked fMRI activity maps (t-values) for three participants for each of the 5 fingers pressing in the extension and flexion directions at 2N. Results were normalized to a surface-based atlas. Maps are shown in the hand-knob region of the left (contralateral) hemisphere. The black dotted line shows the fundus of the central sulcus. The upper inset shows the average sulcal depth.

#### Surface reconstruction and ROI definition

Each participant’s T1-image was used to reconstruct the pial and white-grey matter surfaces using Freesurfer (Fischl, Sereno, & Dale, 1999). Individual surfaces were aligned across participants and spherically registrated to match a template atlas (Fischl, Sereno, Tootell, & Dale, 1999) using a sulcal-depth map and local curvature as minimization criteria. M1 was defined as a single region of interest (ROI) on the group surface using probabilistic cuto-architectonic maps aligned to the template surface (Fischl et al., 2008). We defined M1 as being the surface nodes with the highest probability for Brodmann area 4 and who fell within 1.5cm above and below the hand knob anatomical landmark (Yousry et al., 1997). To avoid cross-contamination between M1 and S1 activities along the central sulcus, voxels with more than 25% of their volume in the grey matter on the opposite side of the central sulcus were excluded.

#### Multivariate fMRI analysis

We used the cross-validated squared Mahalanobis dissimilarity (i.e. crossnobis dissimilarity) to quantify differences between fMRI activity patterns for each pressing condition within each participant (Walther, et al., 2016; Diedrichsen, et al., 2020). Cross-validation ensures the dissimilarity estimates are unbiased, such that if two patterns differ only by measurement noise, the mean of the estimated dissimilarities would be zero. This also means that estimates can sometimes become negative (Diedrichsen, Provost, & Zareamoghaddam, 2016). Therefore, dissimilarities significantly larger than zero indicate that two patterns are reliably distinct.

The fMRI activity patterns were first-level GLM beta-weights for voxels within the M1 ROI mask. Analyses were conducted using functions from the RSA (Nili et al., 2014) and PCM (Diedrichsen, Yokoi, & Arbuckle, 2018) MATLAB toolboxes. The crossnobis dissimilarity *d* between the fMRI activity patterns (***x***) for conditions *i* and *j* was calculated as

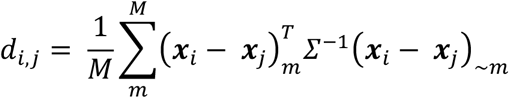

where the activity patterns from run *m* are multiplied with the activity patterns averaged over all runs except *m* (~*m*). *∑* is the voxel-wise noise covariance matrix, estimated from the residuals of the GLM, and slightly regularized to ensure invertibility. Multivariate noise-normalization removes spatially correlated noise and yields generally more reliable dissimilarity estimates (Walther et al., 2016).

The dissimilarities are organized in a representational dissimilarity matrix (RDM). The RDM is a symmetric matrix (number of conditions x number of conditions in size) with off-diagonal values corresponding to the paired distance between two conditions. Values along the diagonal are zero, as there is no difference between a pattern paired with itself.

We calculated an RDM for the matched force conditions separately (i.e. the 1.5N and 2N presses, 10 conditions each), and then averaged the resulting RDMs within each participant. This yeilded one RDM per participant containing the crossnobis dissimilarities between presses of the five fingers in either direction (10 conditions, 45 dissimilarity pairs).

#### Estimating spatial tuning of fingers and direction

We considered the possibility that fingers and directions could be encoded at different spatial scales in M1. We therefore estimated the spatial covariance of tuning for fingers and directions. Within each imaging run, we averaged the fMRI activity patterns (t-values) for each condition across the matched forces (1.5 and 2N). This yielded a vector of 10 activity values per voxel (one value per each finger per direction), which we refer to as an *activity profile*. We modeled the activity profile values (*y*_*i,j*_) of each voxel and partition using three components:

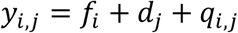

where *f*_*i*_ is the main effect of finger *i*, *d*_*j*_ is the main effect of direction *j*, and *q*_*i,j*_ is the finger x direction interaction effect. We used ordinary least-squares regression to estimate the finger and direction components. The residual from the regression was taken as estimate of the interaction component.

We first reconstructed the activity profiles using only the finger component (*f*), and then estimated the covariance of the finger activity profiles between voxel pairs in M1. These covariances were calculated in a cross-validated fashion: we averaged the reconstructed activity profiles for odd and even runs separately, and then then computed the covariance of the activity profile of different voxels across independent partitions of the data. Given that the estimates for all components contained some noise, normal covariance estimates are biased by the spatial structure of the noise. Cross-validation alleviates the influence of noise on (co-) variance estimation, as the average of the product of noise across odd and even runs is zero.

We then binned the covariances based on the spatial distance between each voxel pair and averaged the covariances within each bin. The first bin included only the cross-partition covariance between each voxel and itself (i.e. the cross-validated estimate of the voxel variances). The second bin contained the covariances between immediately and diagonally neighbouring voxels (1.5 to 2.6mm), the third bin the second layer of direct and diagonally neighbouring voxels (>2.6 to 5.2mm), and so on, up to a total distance of 20.8mm. Finally, we normalized the binned covariances by the cross-validated voxel variances (value of the first bin) to obtain an estimate of the spatial autocorrelation function (ACF) for fingers in M1.

We used the same procedure to estimate the ACF for direction. Importantly, we included both the direction (***d***) and the finger x direction interaction (***q***) components in the activity profile reconstruction. We included the interaction component as we hypothesized that the tuning of voxels to flexion and extension patterns would be different across fingers.

Finally, we estimated the smoothness of the finger and direction ACFs (Diedrichsen, Ridgway, Friston, & Wiestler, 2011). To do this, we fitted a function that decayed exponentially with the square of the distance (*δ*) between voxels (*v*):

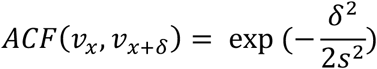

Here, *s* is the standard deviation of the ACF. If neighbouring voxels are relatively independent (i.e. low covariance), the value of *s* will be small. While we can use *s* to express the smoothness of the ACF, the smoothness can also be expressed as the full-width-half-maximum (FWHM) of the Gaussian smoothing kernel that-when applied to spatially independent data- would yield the same ACF. The standard deviation of this Gaussian kernel is 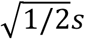, and the FWHM is calculated as:

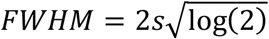

We applied this approach to the reconstructed finger and direction activity profiles separately to estimate the FWHM of fingers and direction M1. The goodness of fit (evaluated with R^2^) of the fitted exponential decays were both high (mean R^2^ of finger ACF=0.960 ±0.008 sem, mean R^2^ of direction ACF=0.908 ±0.020 sem). Although there was a significant difference between the finger and direction model R^2^ (two-sided paired t-test: t_8_=2.412, p=0.0424), the mean difference was quite small (0.052 ±0.021sem).

#### Centre-of-Gravity (CoG) Analysis

We analyzed the activity patterns to determine if there were significant differences in the spatial arrangement of finger flexion and extension, as proposed by Huber et al. (2020). To ensure our analysis closely matched this previous report, we restricted the CoG analysis to include only surface nodes from Brodmann area 4a, as based on the probabilistic atlas (Fischl et al., 2008). We also restricted the analysis to the hand region by selecting only vertices within 1.5cm of the hand knob anatomical landmark. On the flattened activity maps for each finger, we then calculated the centre-of-gravity (CoG) of each map as the average spatial location 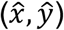 of each surface node (*i*), weighted by its respective *t*-value (*t*):

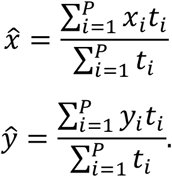

In the above calculations, we set negative *t*-values equal to zero, thereby focusing our spatial analysis on regions that showed activity increases. We used a two-factor repeated-measures MANOVA to test for significant differences between the measured CoGs for different fingers and directions. To summarize the structure of the spatial arrangement, we calculated the pairwise Euclidean distances between the CoG coordinates for each condition, and arranged them into an RDM.

### EMG recording and analysis

#### EMG recordings and preprocessing

In a separate session, we recorded hand and forearm muscle activity to ensure participants performed the task as instructed. During the EMG session, participants were seated upright, whereas during the fMRI session participants lay prone in the scanner. In both sessions, however, we ensured that the arm was in a relaxed position, the palm of the hand was supported by the device, the wrist slightly extended, and the elbow joint slightly bend. Thus, wrist and forearm posture, both known to influence muscle activity during finger movements (Beringer, et al., 2020; Mogk & Keir, 2003) were matched across the two sessions. Participants’ skin was cleaned with rubbing alcohol. Surface EMG of distal muscles of the hand were recorded with self-adhering Ag/AgCl cloth electrodes (H59P-127 repositionable monitoring electrodes, Kendall, Mansfield, Massachusetts, USA). Electrodes were cut and positioned in line with a muscle in a bi-polar configuration with an approximate 1cm inter-electrode distance. Surface EMG of proximal limb muscles were recorded with surface electrodes (Delsys Bagnoli-8 system with DE-2.1 sensors). The contacts were coated with a conductive gel. Ground electrodes were placed on the ulna at the wrist and elbow. The signal from each electrode was sampled at 2000Hz, de-meaned, rectified, and low-pass filtered (fourth order butterworth filter, *f*_*c*_=40Hz).

#### Multivariate EMG analysis

We used the crossnobis dissimilarity to quantify differences between patterns of muscle activities for each movement condition, similar to the fMRI analysis. This metric is invariant to scaling of the EMG signals from each electrode, and has been established in previous work (Ejaz, Hamada, & Diedrichsen, 2015). Briefly, we first calculated the average square-root EMG activity for each electrode and trial by averaging over the press and hold time windows (mean window= 1800ms, up to a max window of 3000ms). We then subtracted the mean value for each electrode across conditions for each run independently to remove any drifts in the signal. These values were then divided by the standard deviation of that electrode across trials and conditions to avoid arbitrary scaling. Finally, we calculated the crossnobis dissimilarity between pairs of EMG activity patterns for different conditions across runs.

### Experimental design of monkey finger individuation task

The behavioural task performed by two male *Macaca mulatta* monkeys (monkeys C and G) has been described previously (Schieber, 1991; Schieber & Rivlis, 2007). Briefly, the monkeys were trained to perform cued single finger flexion and extension presses. Each monkey sat in a primate chair and, similar to the human device described above, their right hand was clamped in a device that separated each finger into a different slot (Fig. 1C). Each slot was comprised of two microswitches (one in the flexion direction and one in the extension direction). One switch was closed by flexing the finger, the other by extending the finger. The absolute degree of movement required to close either switch was minimal (a few millimeters), and therefore the force required to make and hold a successful press was small-similar to the human finger individuation task. Therefore, like the fMRI task behaviour, these finger movements are very close to isometric presses.

A series of LED instructions were presented to the monkey during each trial (Fig. 1D). A successful trial occurred when the monkey pressed the cued finger in the cued direction without closing any other switch. Similar to our human experiment design, the monkeys were trained to hold the cued switch closed for 500ms, before relaxing the finger (Fig. 1D). At the end of a successful trail, the monkey received a water reward. The monkey’s wrist was also clamped in this device, and some trials required the monkey to flex or extend the wrist. Wrist trials were not included in the current analysis. Flexion and extension trials of each finger and wrist were pseudorandomly ordered. In the case of a behavioural error, trials were repeated until successful. Therefore, we excluded all trials with an error and also the successful trials that followed error trials to avoid potential changes in the baseline firing rate of the recorded neuron.

In contrast to the human task, the required force level for the monkeys was the same for all trials – therefore, they did not receive continuous visual feedback about the force produced. Instead, they received small tactile feedback when the switch closed, a feature that was absent from the human task. In spite of these small differences in feedback, the task requirements were well matched across species: Both monkey and humans were required to produce low, well-controlled forces with a single finger, while keeping forces on the non-instructed fingers minimal, either to avoid unwanted switch-closure, or excessive movement of the associated visual feedback.

### Analysis of single cell spiking data

#### Spike rate calculation

Single cells were isolated and spike times were recorded while monkeys performed the finger individuation task. The details of the recordings are reported previously (Poliakov & Schieber, 1999). Each trial was labeled with a series of behavioural markers, indicating the time of trial onset, presentation of condition cue, switch closure, and reward onset. For the spike rate traces plotted in Figure 4, we calculated the spike rate per 10ms bin, aligned to press onset, and smoothed the binned rates with a Gaussian kernel (FWHM=50ms). For the dissimilarity analysis (see below), we calculated the average spike rate over time per trial starting at go cue onset (when the monkey was instructed as to which finger and direction to press) until the end of the hold phase (500ms after switch closure). This time window encompassed a short period of time prior to the start of the finger press and the entire hold duration of the press (Monkey C: mean window= 739ms; Monkey G: mean window=773ms).

#### Multivariate spiking analysis

Similar to the human fMRI and EMG analyses, we computed crossnobis dissimilarities between spiking patterns for different conditions within each monkey. To cross-validate the estimated distances, we restricted our analysis to include cells for which we had at least two successful trials for each finger in both directions. This criteria yielded 44801 trials from 238 cells in monkey C (median number of trials per cell=168, median number of trials per condition per cell=19) and 5535 trials from 45 cells in monkey G (median number of trials per cell=115, median number of trials per condition per cell=12). After calculating the average spike rates, we arranged the spike rates into vectors per condition (Fig. 4B). In order to account for the Poisson-like increase of variability with increasing mean firing rates, we applied the square-root transform to the average firing rates (Yu et al., 2009).

For each cell per condition, we randomly split the square-root spike rates from different trials into one of two partitions. The random splits contained approximately the same number of trials, which ensured that each condition was approximately equally represented in each partition. We then averaged the spike rates within each partition. This yielded two independent sets of spiking patterns per monkey (10 patterns-5 fingers x 2 directions). Per partition, we normalized each neuron’s spike pattern by dividing by the neuron’s max rate across conditions, and then re-weighted the normalized spike rates per cell according to the number of trials per cell (cells with more trials were up-weighted, vice versa for cells with fewer trials). Finally, we calculated pairwise cross-validated Euclidean distances between the two sets of patterns. We repeated this RDM calculation procedure 1000x per monkey, each time using a different random partitioning of the data. We then averaged the RDMs across iterations to yield one RDM estimate per monkey. We note that results were not dependent on the normalization we chose-results were qualitatively consistent when using raw firing rates, z-scoring the firing rates, not applying trial re-weighting, and various combinations of these approaches.

### Kinematic finger model RDM

As in Ejaz et al. (2015), we used the statistics of naturalistic hand movements to predict the relative similarity of single finger representations in M1. In the text we refer to this model as the kinematic model. To construct the kinematic model RDM, we used hand movement statistics from an independent study in which 6 male participants wore a cloth glove imbedded with motion sensory (CyberGlove, Virtual Technologies) while they performed everyday activities (Ingram, Körding, Howard, & Wolpert, 2008). These statistics included the velocities about joint angles specific to each of the five fingers of the participants’ right hands. Positive velocities indicated finger flexion, and negative velocities indicated finger extension.

Because the movement in our finger pressing task was restricted to movements about the metacarpal (MCP) joint of each finger, we used the MCP joint velocities to predict cortical M1 finger similarity. First, we split the data for each joint velocity into two vectors: one for flexion and one for extension, taking the absolute of the velocities in this process. During periods of finger flexion, we set the extension velocity to zero, and vice versa. This resulted in 10 velocity vectors (5 fingers x 2 directions). Then, to account for differences in scaling, we normalized each velocity vector to a length of 1. Finally, we calculated the dissimilarities between pairs of these processed velocity vectors. We averaged these RDMs across the six participants in the natural statistics dataset, yielding one kinematic model RDM.

### Experimental design and statistical analysis

#### Statistical analysis of dissimilarities

We summarized the RDMs by classifying dissimilarities into finger-specific and direction-specific dissimilarities for each participant and dataset. Finger-specific dissimilarities were the dissimilarities between conditions where different fingers were pressed in the same direction (10 pairs for flexion, 10 pairs for extension). Direction-specific dissimilarities were the dissimilarities between conditions where the same finger was pressed in different directions (5 pairs total). Within each category, dissimilarities were averaged. For the human data, we used one-sided, one-sample t-tests to test if mean finger and direction dissimilarities were greater than zero. To compare between the average finger and direction dissimilarities, we used two-sided paired t-tests. We report the mean and standard error of the dissimilarities where appropriate in the text.

#### Statistical analysis of RDM correlations

Pearson’s correlations between the vectorized upper-triangular elements of the RDMs were used to compare different RDMs (Ejaz et al., 2015). To calculate the stability of RDMs, we calculated the Pearson’s correlations between all possible pairs of the participants’ RDMs. This yielded 36 correlations (one per unique participant pair). We Fisher-Z transformed these correlations and calculated the mean and standard error. We used these values to calculate the lower and upper bounds of the 95% confidence interval, assuming normality. Finally, the mean and confidence bounds were transformed back to correlations. We report these values in the text as r=mean [lower bound - upper bound]. The same method was applied to correlations between measured RDMs and model predictions. Note that because we used a within-subject design, the muscle model predictions were specific to each human participant. In contrast, the kinematic model prediction was the same for each participant because data for this model was obtained from an independent study. Paired t-tests were performed on Fisher-z transformed correlations to compare fits between models.

#### Estimating noise ceiling for RDM model fits

Since the dissimilarities between fMRI patterns can only be estimated with noise, even a perfect model fit would not result in a perfect correlation with the RDM of each participant. Therefore, we estimated the *noise ceiling*, which places bounds on the expected model correlations if the model is a perfect fit. We first calculated the average correlation of each participant’s RDM with the group mean RDM (Nili et al., 2014), treating the mean RDM as the perfect model. The resulting average correlation is an overestimate of the best possible fit, as each RDM is correlated with a mean that includes that RDM (and hence also the measurement error of that RDM). To then estimate a lower bound, we calculated the correlation between a participant’s RDM and the group mean RDM in which that individual was removed.

## Results

### M1 fMRI activity patterns differ strongly for different fingers, not for direction

We measured activity patterns evoked in M1 in human participants (n=9) while they performed a near-isometric finger flexion-extension task in a 7T MRI scanner. Participants’ right hands were clamped in a device that had force transducers mounted both above (extension) and below (flexion) each finger (Fig. 1A) to record forces produced at the distal phalanges. The device limited the overall degree of movement to a few millimeters, thereby making the task near-isometric. On each trial, participants were cued to press a single finger in one direction, while keeping the other fingers as relaxed as possible (Fig. 1B). They had to reach the required force level, hold it for 1 second, and then simply relax their hand to let the force passively return to baseline. This aspect of the task instruction was critical to ensure that participants did not activate the antagonist muscles during release.

Figure 2 shows the activity patterns measured in left M1 (contralateral to movement) for three participants during right-handed finger presses at 2N. As previously observed (Ejaz et al., 2015), the activity patterns did not consist of focal regions of activity dedicated to each finger. Rather, the spatial patterns were complex and involved multiple overlapping regions within the M1 hand area. Furthermore, the inter-subject variability in the spatial organization of these patterns was considerable.

One common observation across all participants, however, was that the activity patterns were different between different fingers (e.g. index flexion vs. fourth flexion), but rather similar for flexion and extension of the same finger (e.g. index flexion vs. index extension). We used representational similarity analysis (RSA) to quantify these observations by calculating a measure of dissimilarity (crossnobis dissimilarity, see Methods) between each pair of fMRI patterns. Large dissimilarity values indicate that the two patterns are quite distinct with little overlap. A value of zero indicates that the two patterns are identical and only differ by noise. We restricted the analysis to conditions with matched force levels across flexion and extension. The group-averaged representational dissimilarity matrix (RDM) is shown in figure 3A. Both within the finger flexion and extension conditions, there was a characteristic structure with the thumb activity pattern being the most distinct and neighbouring fingers tending to have more similar activity patterns. Across directions, activity patterns evoked by pressing the same finger in different directions were the most similar. This representational structure was quite stable across participants (average inter-participant Pearson’s r=0.790, 95% CI: [0.754-0.820]).

**Figure 3.**
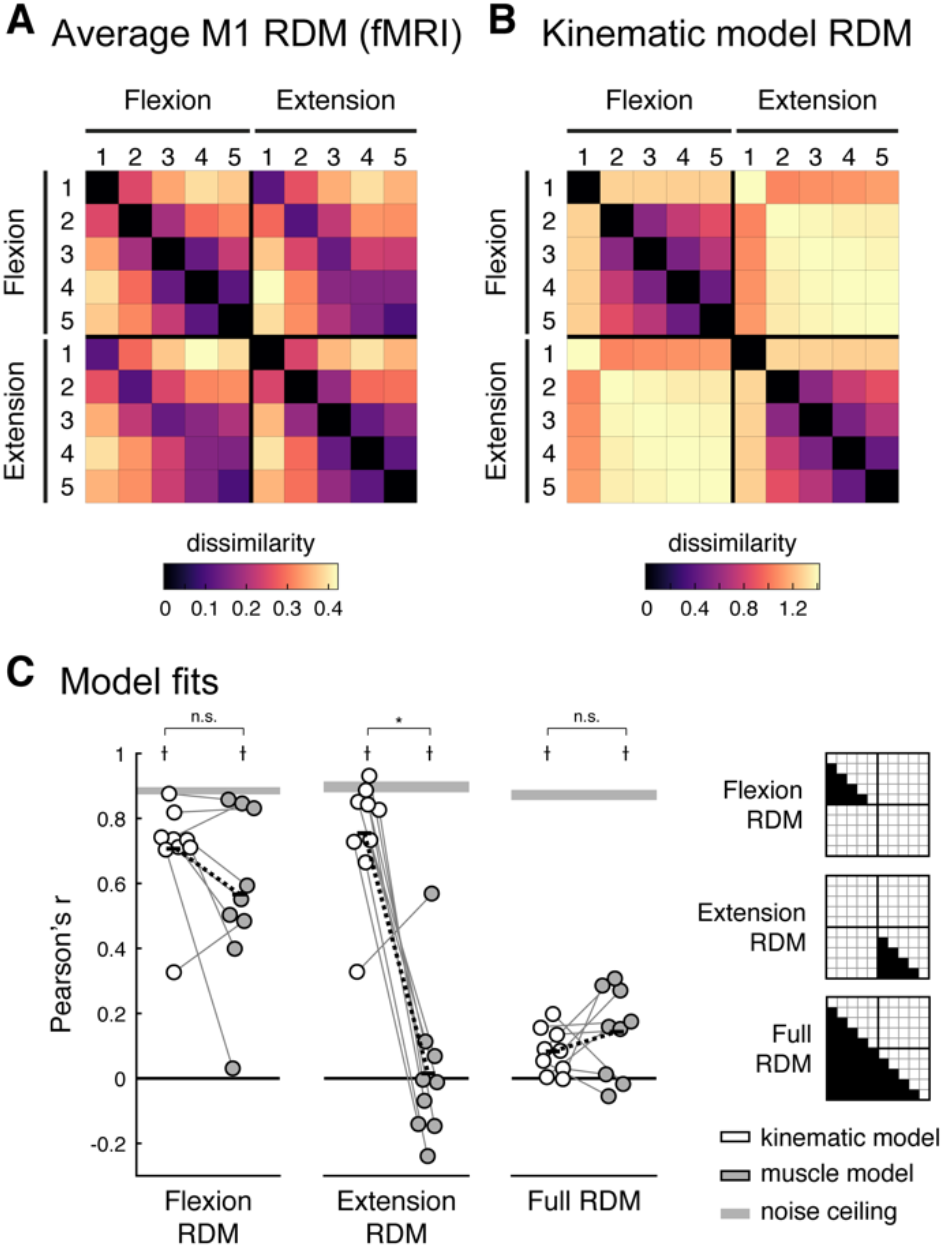
Representational structure of fingers and direction in human M1. (**A**) Group average of the fMRI representational dissimilarity matrix (RDM). (**B**) Predicted RDM from the kinematic model. To aid visual inspection, the values of the RDMs in **A** and **B** are plotted as the square-root of the dissimilarities. All statistical analyses of the RDMs are done on squared distances. (**C**) Model fits (Pearson’s correlation) of the kinematic (white) and muscle (grey) models to the M1 RDM for flexion, extension, and the full RDMs (the indicies for each RDM are shown on the right). The muscle model was specific to each participant and was estimated from the EMG data. The grey bars denote noise ceilings (theoretically the best possible fits). Each dot reflects one participant, and thin grey lines connect fits of each model to the same participant. Black bars denote the means, and black dashed lines denoted the mean paired difference. *significant differences between model fits (one-sided paired t-test, p<0.05); ⟊significantly lower than the noise ceiling (two-sided paired t-test, p<0.05); n.s. not significant (p>0.05).

To obtain predictions for flexion and extension movements, we needed to adapt the natural usage model, proposed by Ejaz et al. (2015). This model used kinematic finger data, specifically the joint-angle velocities of the metacarpal (MCP) joints, recorded while subjects participated in their normal, every-day tasks (data from Ingram et al., 2008). Fingers were predicted to have more similar representations if their movement velocities, across flexion and extension, were positively correlated. For the current experiment, we split the data into periods of finger flexion and finger extension (see methods), resulting in 10 time series, and calculated the correlation between them (after taking the absolute value).

The estimated kinematic RDM (Fig. 3B) showed similar structures within flexion and extension movements. The thumb was the most distinct compared to the other fingers, and for the remaining fingers there was a clear similarity structure with neighbouring fingers more similar than non-neighbouring. This structure very closely mirrored those found for fMRI activity patterns: flexion and extension fMRI RDMs correlated strongly with the corresponding kinematic models for flexion (r=0.727 [0.635-0.800]) and extension (r=0.797 [0.684-0.873]) RDMs (Fig. 3C, white). Compared to the noise ceiling (grey bar in Fig 3C, which reflects the best possible model fit given measurement noise: see methods) the natural use model accounted for 79.9% and 84.9% of the variance in the flexion and extension fMRI RDMs, respectively.

In contrast, the kinematic model completely failed to predict the relationships between activity patterns for flexion and extension. Because flexion and extension of the same finger can never co-occur, the kinematic model predicts that the movements are associated with quite distinct cortical activity patterns. The measured fMRI patterns, however, were rather similar for these two actions. As a result, the full kinematic model was not a good fit to the full fMRI RDM (r=0.086 [0.038-0.133]), much below the noise ceiling (r=0.875 [0.822-0.913]).

Thus, although the statistics of movement co-occurrence was a good predictor for representational similarity between the activity patterns for different fingers (i.e. within flexion or extension), this simple model failed to predict the relative organization of the patterns for flexion and extension of the same finger. Even though flexion and extension of the same finger cannot co-occur, their fMRI activity patterns were highly similar. In the remainder of the paper, we explore a number of possible explanations for this finding and propose a candidate model of the organization.

### Similarities of cortical representations for presses in different directions cannot be explained by the patterns of muscle activity

We first considered the possibility that the structure of similarity between flexion and extension presses can be explained by the patterns of muscle activity required by these movements. Specifically, it is possible that participants co-contracted both agonist and antagonist muscles, or that they activated the antagonistic muscles when returning to baseline. Given the temporally sluggish nature of the blood-oxygen level-dependent (BOLD) signal measured with fMRI, either behaviour could cause the cortical activity patterns evoked during flexion to resemble activity patterns during extension (and vice versa). Therefore, we conducted a control experiment with the same participants outside the MR scanner, during which we recorded surface electromyography (EMG) from 14 sites of the hand and forearm in the participants (Fig. 4A), while they performed the same isometric finger flexion-extension task as in the fMRI session. Performance on the task was comparable to that during the fMRI scan.

**Figure 4.**
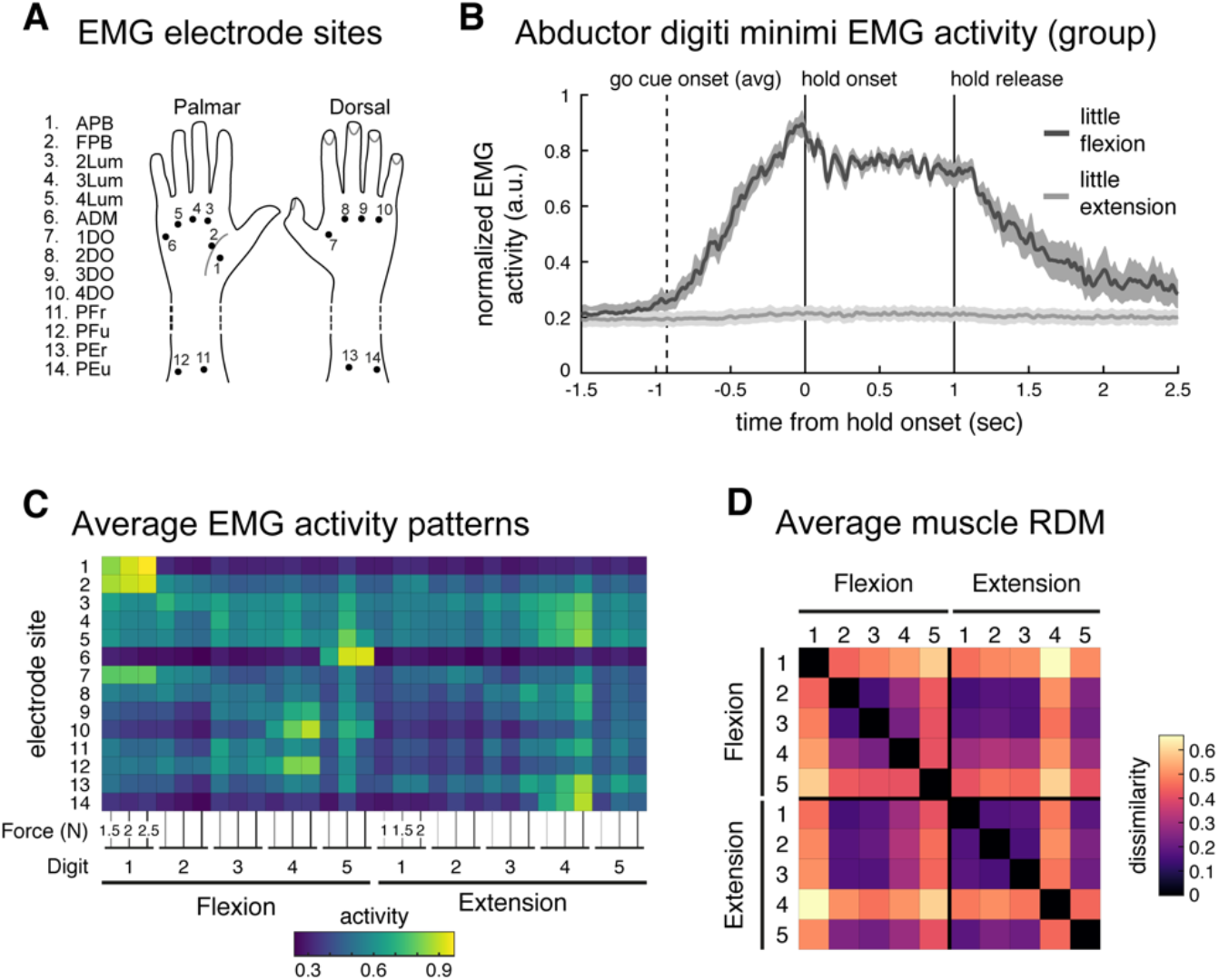
Quantifying similarity of muscle activity patterns during finger flexion and extension. (**A**) Fourteen surface electrode sites. (**B**) Group averaged normalized EMG (normalized, per participant, to peak activity from this electrode across trials) from the abductor digiti minimi (ADM) muscle during 2N little finger (5) flexion (dark grey) and extension (light grey) trials, aligned to hold onset (0s). During extension movement (light grey trace, >1000ms), this flexor muscle was not recruited. Shaded areas reflect standard error of the mean. Traces were smoothed with a gaussian kernel (FWHM=25ms). (**C**) Average muscle activity across participants, normalized by peak activation across conditions (per participant), recorded from the 14 electrode sites during the flexion extension task. Each condition was measured under 3 force conditions. (**D**) Group average representational dissimilarity matrix (RDM) of the muscle activity patterns. As in figure 2, the RDM is plotted as square-root dissimilarities to aid visual inspection.

As an example, the participant-averaged EMG data from an electrode placed above the abductor digiti minimi (ADM) muscle (Fig. 4B) showed that the ADM muscle was recruited only during the flexion of the little finger. During extension of the same finger, the muscle was silent, both during hold and release. In general, we found very little evidence for co-contraction of the antagonist muscle.

For a quantitative analysis, we averaged the muscle activity from the time of the go-cue to the end of the hold phase. The EMG patterns averaged across participants (Fig. 4C) already allow for two observations. First, the muscle activities for the same movement at different force levels were very similar and increased with increasing force. The average correlation across force levels for each finger-direction combination was high, indicating the same muscles were consistently recruited to perform the same finger press across different force levels (within participant correlations: r=0.860 [0.808-0.898]). Second, quite distinct muscle groups were recruited to produce forces with the same finger in different directions. The average correlation between the pattern of muscle activity recruited to press the same finger in different directions was low (within participant correlations: r=0.244 [0.150-0.334]).

We then derived a muscle-based RDM by calculating the crossnobis dissimilarity between normalized activity patterns for each condition. As for the fMRI analysis, we included the patterns for the matched force conditions only. The group averaged matrix RDM (Fig. 4D) was only moderately stable across participants (average inter-participant Pearson’s r=0.480 [0.379-0.570]), likely reflecting the fact that there was some degree of inter-individual variation in electrode placement.

We tested to what degree the patterns of muscle activity, specific to each participant, could explain the cortical similarity structure between individual finger movements within the flexion or extension directions. For the flexion direction, the fit of the muscle model (r=0.611 [0.408-0.757]) was lower than that for the kinematic model in 6 out of 9 participants (Fig. 3C), but the difference did not reach statistical significance (one-sided paired t-test kinematic>muscle: t_8_=1.775, p=0.0569). For the extension direction, the muscle model fit substantially worse (r=0.020 [-0.147-0.187]), significantly less than the kinematic model (one-sided paired t-test kinematic>muscle: t_8_=5.588, p=2.59e-4). This generally confirms the results reported in Ejaz et al. (2015) that the relative similarities of M1 finger flexion activity patterns is better explained by the correlation structure of everyday movements than the correlation structure of the required muscle activity patterns. Our new results now show that this observation generalized also to extension movements.

Critically, however, the muscle activity model did not provide a good explanation for the similarity between flexion and extension patterns. The fit for the full muscle model (r=0.146 [0.055-0.235]) was as poor as for the kinematic model (two-sided paired t-test muscle vs. kinematic: t_8_=1.082, p=0.3108) and significantly below the noise ceiling (two-sided paired t-test noise ceiling vs. muscle: t_8_=12.701, p=1.39e-6). Thus, neither the co-occurrence of movements, nor the pattern of muscle activities can explain the high similarity of activity patterns for finger flexion and extension in M1.

### M1 spiking output differs equally for fingers and direction

To what degree is the high similarity between flexion and extension patterns a function of fMRI as the measurement modality? To approach this question, we analyzed the spiking activity of output neurons in M1 during an equivalent single-finger individuation task in two trained non-human primates (*Macaca mulatta*, data from Schieber & Rivlis, 2005 & 2007). To facilitate this, we had designed the behavioural task for the human fMRI experiment to closely match the task for the monkeys, such that we could make strong comparisons across species and measurement modalities. Figure 5A shows the condition averaged firing rate traces from a single neuron from this data set. This neuron displayed strong preference (increased firing rates) for flexion of the middle finger and extension of the index finger. As previously reported (Schieber & Hibbard, 1993), the population of M1 neurons demonstrated complex, heterogeneous tuning across fingers and directions.

**Figure 5.**
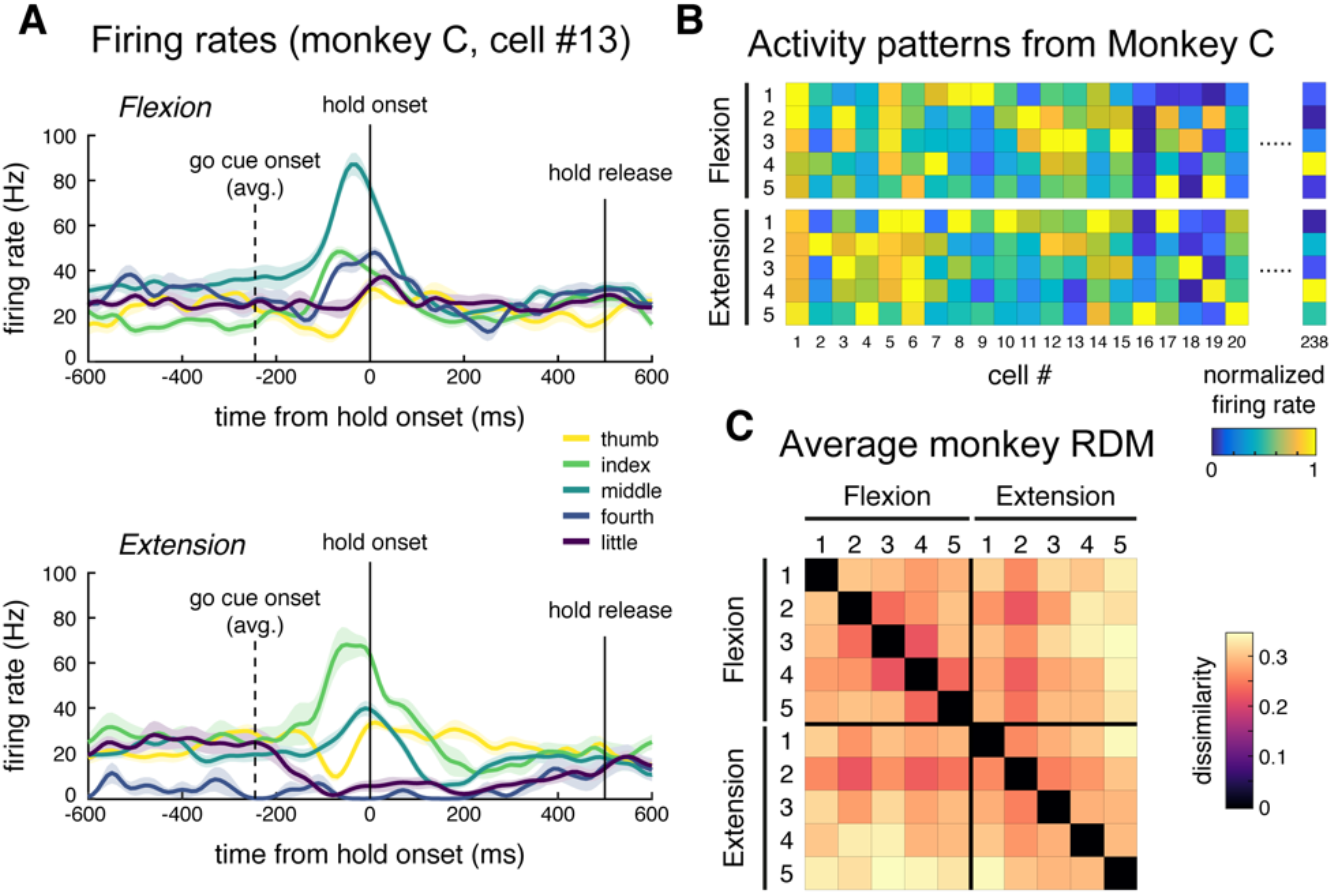
Analysis of M1 spiking activity during monkey single finger flexion and extension. (**A**) Trial averaged firing rates from one cell (monkey C). Traces are aligned to press onset (0s). This cell demonstrates selective tuning to middle finger flexion and index finger extension. Firing rates were calculated for 10ms bins and smoothed with a gaussian kernel (FWHM=50ms). Shaded areas reflect standard error across trials. (**B**) Averaged firing rates for a subset of cells from monkey C, arranged by condition. Cell #13 is plotted in **A**. Firing rates are normalized to the peak rate per cell. (**C**) Average monkey RDM (square-root dissimilarities).

To compare the representational structure from spiking data to that obtained with fMRI, we calculated the mean firing rate for each neuron from the go-cue onset to the end of the hold phase during each trial. We then calculated dissimilarities between the population responses for different conditions (see Methods), similar to the analysis of the human EMG and fMRI data. The average RDM is shown in Figure 5C. Similar to the structure of representations in human M1, the thumb activity patterns for both directions were the most distinct, and neighbouring fingers had more similar activity patterns. In contrast to the fMRI data, however, the spiking patterns for flexion and extension of the same finger were quite distinct.

To quantify this observation, we averaged dissimilarities between different fingers pressing in the same direction (finger-specific) and the same finger pressing in different directions (direction-specific). The finger and direction-specific dissimilarities were close in magnitude for both monkeys (Fig. 6A). Also, the human EMG patterns had roughly matched direction and finger-specific dissimilarities (Fig. 6B). In contrast, the same analysis on the human fMRI data showed a clear and significant difference between these two kinds of dissimilarities (Fig. 6C).

**Figure 6.**
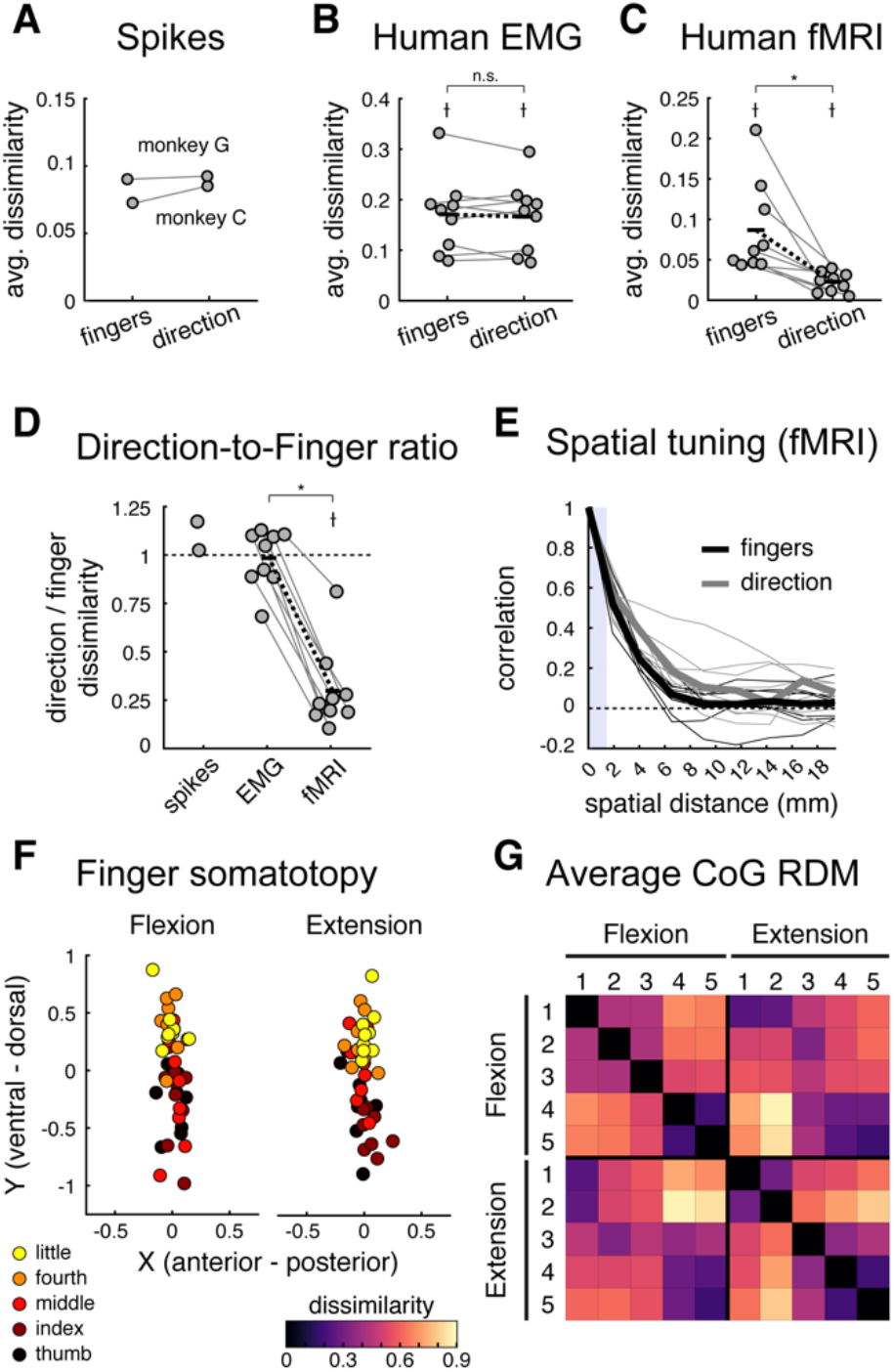
Comparing strength of finger and direction representations across datasets. The average finger and direction-specific dissimilarities for the spiking (**A**), human EMG (**B**), and human fMRI (**C**) datasets. Each dot denotes one participant, and lines connect dots from the same participants. Black bars denote the means, and black dashed lines reflect the mean paired differences. ⟊ dissimilarities significantly larger than zero (one-sided t-test, p<0.05). *significant difference between finger and direction dissimilarities (two-sided paired t-test, p<0.05). (**D**) The ratio of the direction-to-finger dissimilarities for each dataset. Values <1 indicate stronger finger representation. ⟊dissimilatrities significantly lower than one (one-sided t-test, p<0.05). *significant differences between dissimilarity ratios (two-sided paired t-test, p<0.05). (**E**) Estimated spatial autocorrelations of finger (black) and direction (grey) pattern components in human M1, plotted as a function of spatial distance between voxels. No significant difference was observed between finger and direction tuning in M1. The thick lines denote the median spatial autocorrelation functions, and small lines are drawn for each participant for each pattern component. The vertical shaded bar denotes the distance between voxel size, for which correlations can be induced by motion correction. (**F**) Centre-of-gravity (CoG) of activation elicited by single finger presses in the flexion or extension direction for each participant. CoGs were aligned across participants prior to plotting by subtracting the centre of the informative region within each participant (i.e the mean CoG across all conditions). A somatotopic gradient for finger flexion and extension in Brodmann area 4a is visible with the thumb being more ventral and the little finger more dorsal. (**G**) Group average RDM of the paired Euclidean distance between condition CoGs.

For a statistical comparison, we then calculated the ratio between dissimilarities between different directions and dissimilarities between different fingers (Fig. 6D). The fMRI ratio was significantly lower than 1 (mean ratio=0.298 ±0.071; one-sided one-sample t-test: t_8_=-9.858, p=4.72e-6), indicating stronger representation of fingers compared to direction. In contrast, both the spiking patterns (monkey C ratio=1.173, monkey G ratio=1.025) and the human muscle patterns (mean ratio=0.984 ±0.051) differed similarly for different fingers and different directions, with the muscle ratios being significantly larger than those for human fMRI (two-sided paired t-test: t_8_=9.733, p=1.04e-5). Thus, we found a clear difference between the structure of fMRI patterns and the structures of spiking and muscle activity patterns.

We suggest that this difference is informative about the general organization of finger flexion and extension movements in M1. The discrepancy between the two measurement modalities can likely be attributed to two (non-mutually exclusive) differences between fMRI and electrophysiology. First, the fMRI signal is dominated by excitatory inputs and local synaptic signaling, and only partly reflects the spiking activity of output neurons (Logothetis, Pauls, Augath, Trinath, & Oeltermann, 2001). Therefore, the overlapping fMRI activity patterns for flexion and extension might reflect similar inputs and shared local processes within these cortical areas, while the output spiking of these two population remains quite distinct in order to produce the different patterns of muscle activity required for fine finger control.

Second, fMRI samples a proxy of neuronal activity in a coarse manner, averaging across ~200,000 cortical neurons per mm^3^ in M1 (Young, Collins, & Kaas, 2013). Thus, even high-resolution fMRI is biased to functional organization at a coarse spatial scale (Kriegeskorte & Diedrichsen, 2016), and so our results could be caused by an organization where neurons tuned to different movement directions for the same finger (or combination of fingers) are clustered together, while neurons that control different fingers or finger combinations are more spatially separated.

### Spatial organization of finger and direction related fMRI patterns

To investigate the second explanation directly, we attempted to determine whether the activity patterns associated with different fingers were organized on a coarser spatial scale than the patterns associated with flexion and extension of a given finger. Using the fMRI data, we calculated to covariance of the finger-specific and direction-specific activations for each pair of voxels within M1, and binned these covariances according to the spatial distance between voxel pairs (see Methods). If direction is encoded at a finer spatial scale than fingers, we would expect finger effects to be correlated over larger spatial distances.

In contrast to this prediction, the spatial correlation functions for fingers and direction were quite similar (Fig. 6E). We estimated the full-width at half-maximum (FWHM) of the spatial autocorrelation functions. To account for outliers, we evaluated the median FWHMs. The median FWHM of the finger spatial kernel in M1 was 3.22mm (mean=3.44mm ±0.24 sem), comparable to previous reports (Diedrichsen, Ridgway, Friston, & Wiestler, 2011; Wiestler, McGonigle, & Diedrichsen, 2011). The median FWHM of the direction spatial kernel in M1 was 4.65mm (mean=4.77mm ±0.84 sem), and there was no significant difference between the two (two-sided paired Wilcoxon signed-rank test, finger vs. direction: W=11, p=0.2031; two-sided paired t-test finger vs. direction: t_8_=- 1.417, p=0.1942). Therefore, we did not find any direct empirical support for the idea that differences between flexion and extension patterns are organized at a finer spatial scale than differences between fingers. However, our analysis was itself limited by the spatial resolution of 7T fMRI, such that we cannot rule out the possibility that subpopulations for different directions are interdigitated at a sub-voxel scale.

Additionally, we did not find evidence of a substantial spatial separation of flexion vs. extension movements, as was suggested by Huber et al. (2020). These authors observed two sets of digit maps in Brodmann area 4a, with one set being more activated for whole hand grasping, and the other more activated for whole hand retraction movements. From this, the authors suggested that each individual finger map has a preferential function role in guiding flexion and extension movements. To test this idea with our fMRI data, we calculated the centre-of-gravity (CoG) of the activity maps for each finger pressing in the flexion and extension directions in Brodmann area 4a (see Methods).

As shown in figure 6F, both finger flexion and extension CoGs revealed the expected overall somatotopic gradient, with thumb movements activating more ventrolateral areas and the little finger activating more dorsomedial areas in 4a (2-factor repeated-measures MANOVA, finger factor: Wilks’ Λ_(4,32)_=0.28, p=2.2075e-6). However, there was no significant difference in these digit maps across flexion and extension movements (2-factor repeated-measures MANOVA, direction factor: Wilks’ Λ_(1,8)_=0.88, p=0.6427; finger x direction interaction: Wilks’ Λ_(4,32)_=0.65, p=0.0793). We then calculated the pairwise Euclidean distances between the condition CoGs (Fig. 6G) and compared the between and within finger distances, as done previously. Replicating the results from the fMRI RSA analysis, we found that pressing different fingers resulted in more spatially distinct activation patterns compared to pressing the same finger in different directions (mean ratio=0.67 ±0.04; one-sided one-sample t-test ratio<1: t_8_=-8.003, p=4.356e-5). This finding in inconsistent with the idea of separate flexion and extension finger maps.

## Discussion

Here we investigated how the population activity in M1 is organized for control of flexion and extension of single fingers. We analyzed M1 population activity measured in humans with 7T fMRI and spiking data from NHPs while participants made isometric single finger presses in either direction. Importantly, we ensured the behavioural tasks in both experiments were carefully matched to allow us to compare results across the two datasets.

We first demonstrated that the representational structure of single finger flexion or extension presses in human M1 measured with fMRI were relatively well explained by the natural statistics of every-day movements, replicating the flexion results reported in Ejaz et al. (2015) and extending them to single finger extension movements. The same model, however, failed to correctly predict the relationship *between* flexion and extension movements. Because flexion and extension of the same finger cannot temporally co-occur, the model predicted quite separate representations for the two actions. In our data, however, we observed the opposite effect – cortical M1 activity patterns measured with fMRI in humans were very similar for the flexion and extension of the same finger, as compared to the quite distinct patterns for different fingers. We also analyzed spiking data from a similar task in two monkeys and found that the similarity of finger flexion and extension were specific to fMRI: In the monkey electrophysiological recordings, different movement directions were associated with distinct patterns of neuronal activity.

The discrepancy between the fMRI and electrophysiological measures suggest a specific organization of finger flexion and extension movements in M1 (Fig. 7). This suggested architecture has two characteristics that likely contribute to the observed difference between measurement modalities.

**Figure 7.**
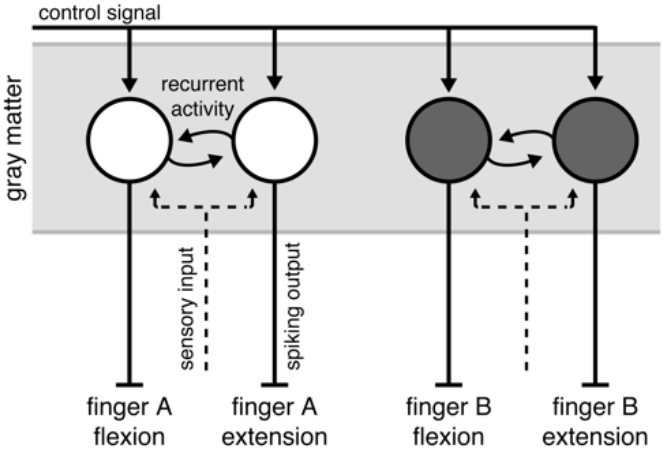
Summary model of M1 organization. Output neurons in M1 produce complex patterns of muscular activity. We refer to groups of neurons that, together, evoke a complex pattern of muscle activty that results in single finger movements as functional units (circles). These functional units receive a control signal input for the upcoming movement (solid lines with arrows). Functional units that evoke movements of the same finger in opposite directions receive common inputs (dashed lines) and share strong recurrent connections (circular lines). The spiking output (solid lines without arrows) of these units, however, is directionally specific. Additionally, under the spatial scale model, functional units tuned to finger movements in different directions are clustered together according to their finger tuning.

First, we hypothesize that neurons that contribute to the flexion of a finger receive similar sensory input as neurons that contribute to the extension of the same finger (dashed line, Fig. 7). There is evidence in the literature to support such an organization. In macaque M1, single neurons tuned to torque production at the shoulder integrate information from the shoulder and elbow joints to facilitate rapid corrective responses to mechanical arm perturbations (Pruszynski et al., 2011). Thus, these neurons receive common sensory input about the shoulder and elbow joints, but the output is largely specific to movements about the shoulder. Additionally, units controlling flexion and extension of the same finger a likely to directly communicate with each other (curved solid arrows, Fig. 7). Such coordination would be necessary to orchestrate fast alternation of finger movements and to finely control the grip force during object manipulation.

This organization would lead to highly similar fMRI activity patterns. In cortical grey matter, the BOLD signal measured with fMRI reflects mainly excitatory postsynaptic potentials (EPSPs), caused by input to a region or recurrent activity within a region (Logothetis et al., 2001). This is because much of the metabolic costs associated with signal transmission arise from re-establishing resting membrane potential of neurons after an EPSP (Attwell & Laughlin, 2001; Magistretti & Allaman, 2015; Yu et al., 2018). Given that the input to subpopulations controlling flexion and extension of the same finger will be highly temporally correlated, the fMRI activity patterns for the two movements should also be very similar.

At the same time, the two subpopulations need to produce distinct spiking outputs. To do so, the populations must receive a control signal input that defines whether to flex or extend a finger. Indeed, in our fMRI data, although flexion and extension patterns for the same finger were highly similar, we could still discriminate between the patterns. This control signal would influence how neurons react to sensory inputs and the information they exchange. Thus, the observed local variations in metabolic activity would be dissociated from the local neural firing rates (Picard, Matsuzaka, & Strick, 2013).

As a second characteristic, we also hypothesize that units controlling muscle patterns that produce flexion and extension of the same effector are spatially co-localized to support fast and efficient communication. Because fMRI samples activity in a coarse manner, even high-resolution fMRI is biased to functional organization at a coarse spatial scale (Kriegeskorte & Diedrichsen, 2016). Therefore, features that exist at fine spatial scales in the neural population are under-represented in fMRI activity patterns. Our results could therefore be caused by an organization where neurons tuned to different movement directions for the same finger (or combination of fingers) are clustered together, while neurons that control different fingers or finger combinations are more spatially separated. We did not find any evidence for a difference in spatial organization of fingers and direction in the fMRI data. However, given that this comparison itself is limited by the spatial resolution of fMRI, we cannot rule out that differences in the fine-grained spatial organization also contributed to the observed effect.

Although we experimentally studied the flexion and extension of single fingers, we do not suggest that isolated finger movements are explicitly represented in M1. Rather, M1 output neurons will produce a complex pattern of muscle activity. This complexity likely arises because the neuronal populations are optimized to produce muscle activities which elicit combinations of finger movements that are useful in everyday tasks (Poliakov & Schieber, 1999; Gentner & Classen, 2006; Ejaz et al., 2015). When we measure activity patterns related to movements of isolated fingers, we simply observe the specific combination of neuronal populations that need to be active to move a single finger (Schieber, 1990). The core of our hypothesis is that populations of neurons that produce opposing muscular patterns form a functional unit with increased communication, common sensory input, and potentially also spatial co-localization.

Our findings are at odds with the organization suggested by Huber et al. (2020). Using high-resolution functional imaging in humans, the authors reported evidence of two spatially distinct finger maps in M1, one for flexion and one for extension. Consistent with Huber et al., we found that individuated finger activity patterns in M1 are fractured and have multiple hotspots (Fig. 2). However, we found no evidence for a clear spatial separation of flexion and extension finger into two action maps (Fig 6F-G). Even though the spatial resolution of BOLD imaging in our study was lower than that of the blood-volume based method employed by Huber et al., we should have been able to detect larger spatial separations between flexion and extension movements than between individual fingers. Instead, the opposite was the case. Both the RSA and the spatial analyses showed greater differences between fingers than between directions. These results, however, are not unexpected. Partial inactivation of neurons in the hand area of macaque M1 result in a complex loss of flexion and/or extension movements of different fingers (Schieber & Poliakov, 1998), and electrophysiological recordings from this same area show flexion and extension preference is not spatially clustered (Schieber & Hibbard, 1993). We believe that the differences between our results and those of Huber et al. are likely explained by the fact that Huber et al. did not study flexion and extension of individual fingers, but relied on a large spatial gradient detected between whole-hand grasping and retraction. We think this is problematic, as the control requirements of individual finger movements is qualitatively different from those of whole hand grasping. That is, neuronal activity during whole hand grasping is not the sum of the neural activity during individuated finger flexion movements (Ejaz et al., 2015), but rather engages a different control mechanism. Consistent with this idea, electrophysiological studies have shown that the neural control of whole hand and individuated finger movements relies on different neural subpopulations (Muir & Lemon, 1983; Lemon, 2008).

There are of course many caveats when comparing results across different recording methodologies, experimental setups, and species. While we tried to make the behavioural tasks across human and macaques as similar as possible, species differences or the extensive training for the non-human primates may account for some of the differences.

Overall, however, we believe that the comparison between fMRI and spiking provides some interesting insights into the organization of the hand region of the primary motor cortex. Cortical representations of single finger movements are not purely dictated by the kinematics of hand usage. We posit that the deviation from this organization appears to reflect a control process, where neurons tuned to movements of a specific finger receive common sensory input and share local recurrent processes. These tightly coordinated populations then produce the spiking output that needs to be quite distinct for the flexion and extension of the same finger.

## Acknowledgements

The work was supported by Canada First Research Excellence Fund (BrainsCAN) collaborative postdoctoral grant awarded to NE, JW, and SA, and a Discovery Grant from the Natural Sciences and Engineering Research Council (NSERC, RGPIN-2016-04890) to JD. Functional imaging costs were partly supported by a Platform Support Grant from Brain Canada and BrainsCAN. SA and EK are supported by doctoral scholarships from NSERC (PGSD3-519263-2018 and CGSD3-519372-2018, respectively). JW is supported by a BrainsCAN Postdoctoral Fellowship. MHS was supported by NINDS grants NS27686 and NS102343. JP is supported by the Canada Research Chairs program. We thank Marcus Saikaley for help with human EMG data collection.

